# Predation, community asynchrony, and metacommunity stability in cyanobacterial mats

**DOI:** 10.1101/2022.10.07.511315

**Authors:** Ethan C. Cissell, Sophie J. McCoy

## Abstract

The dynamism of ecological interactions in rapidly changing ecosystems can be understood only by linking community context to population dynamics. Holistic characterization of such mechanisms requires integrating patterns of variability across scales. Here, we integrated observational, experimental, and theoretical approaches to unify local and regional ecological processes driving the dynamics of benthic cyanobacterial mats on coral reefs off the island of Bonaire, Caribbean Netherlands. Community and metacommunity dynamics of mats were tracked for 49 days alongside quantification of macropredation pressure from fishes. We tested the hypothesis that enhanced predation would result in decreased mat persistence *in situ*. Finally, we constructed a cellular automaton model to predict patterns in mat metacommunity dynamics across different scenarios of top-down and bottom-up control and dispersal. Cyanobacterial mat metacommunities were temporally stable across the study, stabilized by asynchrony in the dynamics of communities. Diverse reef fishes foraged on mats *in situ* and experimental increases in predation pressure decreased the instantaneous mortality rate of mat communities over mat communities experiencing natural levels of predation pressure. Theoretical simulations suggested that dispersal conveys a rescuing effect on mat metacommunity abundance under scenarios of strong trophic control.

## INTRODUCTION

Mechanisms that drive changes in the abundance of organisms operate across many distinct spatiotemporal scales, and with variable frequencies and magnitudes (Connell et al. 1997), requiring integrative study to deal with this issue of variability across scale (Levin 1992). Theoretical developments in metacommunity ecology provide a robust framework for distilling the high dimensionality of complex species interaction networks - including all indirect and direct network linkages among populations - across local and regional scales into meaningful axes describing spatial stability vs. dynamism (Leibold et al. 2004). Global change and local stressors can modify interaction networks among competitors (McCoy and Pfister 2014) and trophically linked populations (Petchey et al. 1999, Falkenberg et al. 2013). Often, this elicits trophic guild-specific revisions to the predominance of competing trophic and competitive control mechanisms, especially in microbial systems, which makes it difficult to generalize traditional paradigms of top-down control and competitive network structure in multitrophic microbial assemblages in the current Anthropocene (Ethan C. Cissell and McCoy 2022). The effects of shifting species linkages are further complicated by hierarchical interactive effects of interspecific competition and predation in structuring prey populations (Paine 1966), as well as differential responses to changing environmental contexts among trophic interactors (McCoy and Kamenos 2018). Collectively, this necessitates an explicit understanding of individual responses (McCoy et al. 2018), and all pairwise and interactive linkages (including indirect links such as interaction modifications [e.x., Wootton 1993]), to fully characterize mechanisms that connect abiotic and biotic context with patterns in population and community persistence (Hunter and Price 1992, Power 1992). Understanding how trophic links affect population dynamics and community structure via both direct and indirect effects is especially critical for effective intervention and control of nuisance organisms, such as blooms of toxic microorganisms (often dominated by algal taxa), which are dramatically increasing in abundance across aquatic ecosystems in response to Anthropogenically-mediated environmental change (Huisman et al. 2018). While planktonic cyanobacterial and eukaryotic algal blooms receive significant research and public attention, benthic blooms are underappreciated despite inhabiting and impacting some of the most threatened ecosystems globally.

The influence of both local and global stressors is driving near-ubiquitous increases in the proportional cover of conspicuous benthic cyanobacterial mats on coral reefs worldwide (reviewed in Ford et al. 2018), including increasing the relative cover of spatially discrete, horizontally-spreading mat carpets (de Bakker et al. 2017) and vertically tufting cyanobacteria (Ford et al. 2021), as well as increasing the duration and magnitude of extensive (i.e., kilometer-scale) benthic bloom events (Hewson et al. 2001, Paul et al. 2005). Similar to microbial mats found in other systems (Bolhuis et al. 2014), cyanobacterial mats found on coral reefs are complex and strongly cooperative microbial consortia (Cissell and McCoy 2021, Stuij et al. 2022). Coral reef cyanobacterial mats, while structurally built by phototrophic cyanobacterial taxa, also contain a taxonomically and functionally diverse suite of microbial eukaryotes, heterotrophic bacteria, and viruses (Cissell and McCoy 2021, Stuij et al. 2022). Expanding cover of cyanobacterial mats generates complex feedback loops with other major causes of reef degradation, and may exacerbate the velocity and magnitude of existing trajectories of reef decline (Ethan C. Cissell and McCoy 2022). For example, expanding cyanobacterial mat cover may increase the transmission of coral diseases, further contributing to the erosion of coral cover and leading to the eventual loss of topographic complexity (Cissell et al. 2022). A better understanding of the mechanisms promoting cyanobacterial mat cover on reefs and controlling mat dynamics across spatial scales is an increasingly urgent challenge for coral reef ecologists and managers in the current Anthropocene (Cissell et al. 2022, Ethan C. Cissell and McCoy 2022).

The focus of most research on coral reef benthic cyanobacterial mats has been on mat formation and growth. Mat growth dynamics have been continuously linked to bottom-up processes (i.e. release from bottom-up control via nutrient enrichment and warming) in driving the creation of cyanobacterial mats (Kuffner and Paul 2001, Albert et al. 2005, Brocke et al. 2015a). Little focus, however, has been given to the dynamics of mature (post-formation) cyanobacterial mat communities on coral reefs, and so the relative influence of top-down trophic interactions on mat community dynamics following mat establishment is largely unknown (Ethan C. Cissell and McCoy 2022). Macrofaunal predation is generally considered to be a strong biotic filter preventing the establishment of microbial mats (Fenchel 1998). Microbial mats, then, generally only form in extreme environments that limit pressure from top-down forcing via physical exclusion of predatory macrobiota (Bolhuis et al. 2014). This logically suggests that top-down trophic interactions may strongly influence coral reef cyanobacterial mat bloom dynamics. Predatory interactions with both reef fishes and viruses have been previously documented with coral reef benthic cyanobacterial mats (Hewson et al. 2001, Cissell et al. 2019, Cissell and McCoy 2021, Ethan C Cissell and McCoy 2022), and correlative assessments of fish density and site-scale mat cover suggest some influence of predation on mat dynamics (Reverter et al. 2020). The spatiotemporal scales at which processes in this experimental system proceed allow dynamics across multiple scales of ecological organization, from populations to metacommunities, to be readily observable and amenable to experimental manipulation. Mat community patches on the reefs surrounding the Caribbean island of Bonaire form spatially distinct mats with discrete patch boundaries, which may be longitudinally tracked macroscopically for evidence of predation and for broad community dynamics while efficiently solving the classic issue of community delineation and circumscription (Crowley 1978). Furthermore, recent molecular evidence from the comparison of average total nucleotide identity of Metagenome Assembled Genomes (MAGs) reconstructed among multiple spatially distinct benthic cyanobacterial mats on the fringing reefs of Bonaire revealed strong genomic conservation among mat communities, and suggests that cyanobacterial mats form true, dispersal-linked metacommunities on reefs (Cissell & McCoy *unpublished manuscript*).

Here, we integrate observational and field experimental characterizations of local trophic interactions with documentation of spatial dynamics at multiple spatial scales and model simulations to understand and unify local and regional ecological processes driving the dynamics of benthic cyanobacterial mats on coral reefs off the island of Bonaire, Caribbean Netherlands. Natural patch (community) and metacommunity dynamics of cyanobacterial mats were tracked photographically alongside observational documentation of predation pressure from fishes. Further, we employed an *in-situ* coring experiment (simulated wounding) to test the hypothesis that enhanced predation would result in decreased mat patch persistence via a strong community response to top-down pressure (Fenchel 1998). We constructed a cellular automaton model to predict patterns in mat metacommunity dynamics across different scenarios of top-down and bottom-up control and dispersal. Finally, we curated a selection of relevant natural history observations made during our study period that further contextualize the patterns and processes presented herein (presented in Supplementary Information: Supplementary Discussion). By pairing a longitudinal characterization of the structural state of mat landscapes and interaction network linkages with experimental manipulations of predation and theoretical simulation, we can begin to contextualize and link patterns observed in space and time to the trophic processes generating structural dynamics across scales of ecological organization in cyanobacterial mats.

## MATERIALS & METHODS

### Tracks of natural and experimentally cored cyanobacterial mat communities, and assessment of metacommunity dynamics

Benthic cyanobacterial mats were tracked along a stretch of reef (146.2m straight-line length) on the southern leeward side of Bonaire, Caribbean Netherlands, in the Caribbean Sea (surveilled reef stretch extended from N12° 06.083’ W68 17.269’ to N12° 06.201 W68 17.328’). On 28 unevenly distributed, unique sampling days spanning a total duration of 47 days (23 May 2019 – 09 July 2019), we took repeat, scaled photographs of 26 mats of the dominant orange-red morphotype (Figure S1) using an integrated-camera PVC monopod (1m height). To establish which mats would be targeted for repeat sampling, mats growing on sediment in a ± 4m depth band (10 - 18m depth) were marked along a transect at 14m depth parallel to the reef slope. Although mats are also ubiquitous on hard substrate and living benthos (Ritson-Williams et al. 2005, Cissell et al. 2022; Figure S2), only mats growing on sediment were marked to gain a better baseline understanding of mat bloom dynamics in the absence of any influence of competition/facilitation from close association with macro-benthic organisms. Individual mat communities were defined as spatially distinct mat individuals with discrete patch boundaries from other entire mats (Figure S1). Generally, several meters of reef separated individual mat communities. Mats were semi-permanently marked with two plastic stakes and a PVC place-marker to allow for repeat monopod placement over the 49 days and to facilitate intra-image georeferencing in later analyses.

To explore the effects of disturbance from top-down forcing on mat dynamics, an additional 27 mats were identified as described above and a 1.3cm diameter core was experimentally removed from both the interior of the mat and from the mat’s border using a PVC corer and forceps to simulate predation from reef fishes (core size predetermined from mean size of bite scarring measured in mat images taken in Jan 2019; Cissell, Pers. Obsv.). These mats were marked and repeatedly photographed as described above.

Dynamics in the site-scale cover of sediment-bound cyanobacterial mats was used as a proxy for assessing mat metacommunity dynamics at our study site. Mat cover across a depth range of 7m – 19.8m across the length of the study site from 5 separate sampling dates was assessed in 204 haphazardly placed photoquadrats (0.25m^2^) from manual annotation of 50 randomly placed points per quadrat in *CoralNet* beta. Only cyanobacterial mats growing on sediment were annotated as cyanobacterial mat in this analysis – mats overgrowing hard substrate were annotated as EAM. Because these functional groupings were intentionally designed differently from conventional grouping structures to capture and track cyanobacterial mat dynamics specifically on sediment, we make no inference on the cover of other benthic groupings and dissuade use of these benthic cover data toward alternative purposes to those stated herein.

### Surveys of macropredatory pressure

Grazing rates of reef fishes were quantified using diver-independent, fixed-point, timed behavioral observations using either a Nikon Coolpix W300 or a GoPro Hero4 equipped with a red filter. Observations were made on 22 individual mat communities across 5 sites found between 13.2m and 17.3m depth (mean 14.9m) between the hours of 10:00 and 16:00 corresponding to the peak feeding time of most grazing fish species in the Caribbean (Bruggemann et al. 1994b) for a mean observational duration of 26 minutes per mat. Survey cameras were placed on reef substrate away from the focal mat such that the entire mat area was visible and surveilled within the frame. The first minute of each behavioral observation period was removed from analysis to allow the behavior of the fish community to acclimate to camera presence. During video analysis, each bite that visibly removed biomass was counted, and the identity of the species biting the mat was recorded. Foraging by spotted goatfish (*Pseudupeneus maculatus*) and yellow goatfish (*Mulloidichthys martinicus*) individuals mostly consisted of the following feeding modes: shovel, push, and skim surface (Krajewski et al. 2006). Though these goatfish individuals were likely not foraging directly on cyanobacterial mat biomass, their foraging activity disturbs mat biomass similarly to predation from other reef fishes actively grazing cyanobacterial mat biomass, and so data on their foraging are presented as ‘bites’, where each individual foray was counted as a single ‘bite’ regardless of duration or magnitude.

### Cellular automaton model overview, parameterization, and simulation procedure

Model simulations using C^++^ were used to predict patterns in mat metacommunity dynamics across gradients of top-down and bottom-up control and dispersal, motivated by the need for a mechanistic understanding of the decoupling between nutrient control and management outcomes for standing mat cover on reefs (model code provided in Github link for reviewers). The model presented is based on the cellular automaton model previously presented in McCoy et al. (2016). The simulated site was represented by a grid of 5,000 × 5,000 cells, with each grid cell corresponding to an area of reef 50mm^2^ in size (roughly corresponding to average bite sizes measure in Jan. 2019; Cissell, Pers. Obsv.) for a total modeled area of 1,250m^2^. Opposing edges in the model grid were connected to wrap the grid into a torus to avoid edge effects (McCoy et al. 2016). The bare grid represents colonizable reef substrate. Coral reef cyanobacterial mats have been observed overgrowing most possible benthic substrates, both biotic and abiotic (Figure S2), and therefore no distinction is made among benthic substrates to match the assumption of equal occupational probability across substrate types. A single model time step is equivalent to a period of 24hrs, and each model was run for a total of 365 time steps for a total duration of a 1-year time period per simulation.

The starting simulation grid matrix was generated in QGIS ver. 3.16 (Hannover). A raster layer of bare substrate was randomly populated with cyanobacterial cells following a binomial distribution with an N of 1 and a probability of 0.2, creating a starting site-scale abundance of 20% cyanobacterial mat. This generated grid was input as the starting grid for every model run.

A total of 64 unique parameterizations were simulated, including all possible permutations of 4 levels per parameter across 3 unique parameters. Parameters that were varied in this model were mat DISPERSAL, mat GROWTH RATE, and DISTURBANCE (broadly encompassing macropredation, micropredation, and goatfish disturbance each with independent probabilities). Levels of each parameter were broadly classified into the following bins: Zero, Low, Medium, and High. Parameterizations correspond to a probability of each event occurring within a model time step. Exact numerical probabilities for each parameter are provided in Table S1. For each model ‘decision’, a random number from 0-1 was pulled from a random number generator and compared to the parameterized probability of each event to determine the outcome of the event. Within a time step, focal cells were chosen at random. Focal cells that were not occupied by cyanobacterial mat could have mat successfully dispersed into them with probability DISPERSAL, filling the cell with mat when successful. It should be noted here that the process of dispersal is modeled as a generative colonization event, and not necessarily as population connectivity among spatially distinct mat communities. Focal cells that were instead occupied with mat and that had an empty neighboring cell (pulled at random) could then grow into that empty neighboring cell based on GROWTH RATE. Each focal cell could only grow into a single empty neighboring cell per time step (maximally one growth event per focal cell per model step). Following this growth step, focal cells that were occupied were then subject to a possibility of disturbance from goatfish, macropredation (from consumptive reef fishes), and micropredation based on each respective probability from DISTURBANCE. Either when a focal cell had been disturbed (became empty substrate) or passed all disturbance events without success (remained mat), a new focal cell was chosen (without replacement) and the loop continued. Only one disturbance event per focal cell per time point was possible. Total counts of bare substrate and mat cells were tallied at the end of each time step. Model outputs (abundances from counts/total) are presented as means and standard deviations from 100 independent model runs per parameterization combination.

### Statistical procedures

All statistical analyses were conducted using the R programming language (ver. 3.6.2) implemented in RStudio (ver. 1.2.5033). Data visualization was performed using *R::ggplot2* (ver. 3.3.3). Model assumptions for all statistical models (across all chosen likelihoods; unless otherwise specified) were assessed from model residuals graphically using *R::DHARMa* (ver. 0.3.3.0).

To assess metacommunity dynamics at the study site across the study duration, the site-scale cover of cyanobacterial mats was modeled against both benthic sampling date and depth of photoquadrat (n=204) as predictors using linear models (Gaussian likelihood). The distribution of photoquadrats across depths across sampling dates was initially assessed graphically (Figure S3). A nested model structure was fit including an additive or interactive effect of both predictors. The interactive terms were not significant (all pairwise p > 0.05; overall term F = 1.29, df = 4, p = 0.27), and a likelihood ratio test - conducted to determine if adding the complexity of the interaction term improved model fit - suggested that the interaction term could be dropped (p = 0.27). Parameter estimates and error structures presented are from the additive model. Post-hoc pairwise Wilcoxon Rank Sum Tests with Bonferroni p-value corrections were used to assess pairwise differences among levels of sampling date.

To test for an effect of the experimental coring treatment on the overall probability of a mat (n=53) dying during the study period, a binomial Generalized Linear Model (GLM) was fit, where the response variable was a 0 or 1 corresponding to if a mat died during the study period (“0”) or not (“1”). Both Treatment and Start Date (i.e., the date of tracking onset) and their interaction were included as predictors in the model. A Kruskal-Wallis test was used to test for a significant difference in start dates among treatments. No significant difference in start date among treatments was detected (K-Wχ^2^ = 0.02, df = 1, p = 0.88), however a likelihood ratio test between the fully interactive model and a model fit without Start Date as a predictor suggested that Start Date should be included in the analyzed model (p=0.02). The significance of fixed effects in each model, including that of the interaction term, were assessed using Type II Wald’s χ^2^ tests conducted using car [3.10-10]::Anova. The interaction term between Treatment and Start Date in this interactive GLM was not significant (df = 2, F = 1.63, p = 0.21), offering further verification that there was no significant difference in start dates between treatments. The instantaneous mortality rates of mats in each treatment of the field simulated predation experiment were predicted by fitting exponential decay curves (logistic regression) to the experimental data. A quasibinomial GLM was fit to the binomial response variable of mat death (structure described above) against observation duration in days without intercepts. The model was fit using the quasibinomial family distribution with a log link function, with separate parameters estimated per treatment. The significance of model terms was assessed using a χ^2^ test.

To model bite count on cyanobacterial mats, we fit a generalized linear mixed effects model using glmmTMB::glmmTMB (ver. 1.0.2.1) with a negative binomial distribution and a log link, treating site and fish species as fixed predictor variables and allowing the intercept to vary by mat identity nested within site, including an offset of the log of video duration (in seconds). The initial phase and terminal phase of species, where available and applicable, were fit as separate levels of the predictor Species and were not separately included as a Phase predictor variable because different phases were not observed and not applicable for all species observed consuming cyanobacterial mat biomass. A Type II Wald’s χ^2^ test was used to assess the significance of fixed effects parameter estimates. Bites taken by bicolor damselfish were removed from this model because the direct effects from their bites are small in magnitude (minimal biomass removal).

## RESULTS

### Cyanobacterial mat metacommunities are temporally stable, with asynchronous instability of component communities

The dynamics of spatially distinct individual benthic cyanobacterial mat communities and the broader site-scale metacommunity were photographically monitored for 49 days along a stretch of reef 146.2m long in straight-line length on the leeward fringing reefs of Bonaire. The metacommunity at this site was temporally stable across the study, slightly increasing after initial sampling to a consistent cover (Fig. 5.1a; LM; df = 4; F = 4.79; p = 0.001; PWRST p=0.02, 0.0002, 0.042 between initial and 2^nd^, 3^rd^, and 4^th^ sampling, all other p > 0.05). Mean metacommunity cover across the study duration was 19.6% ± 4.94% SD (Fig. 1a). Site-scale cyanobacterial mat cover showed a significant trend with depth, increasing by ∼1.2% benthic cover per meter depth increase (LM; y = 0.012x - 0.04; df = 1; F = 9.42; p = 0.002; Figure S4). High abundances of cyanobacterial mats have been reported at mesophotic depths in other reef ecosystems (Sellanes et al. 2021) and on the mesophotic reefs around Bonaire (van Heuzen 2015). Additionally, this study site is marked by the presence of a sand channel at deeper depths connecting a double reef structure whose flanking regions provide a protected habitat for cyanobacterial mat proliferation, making this trend with depth unsurprising.

**Figure 1.**
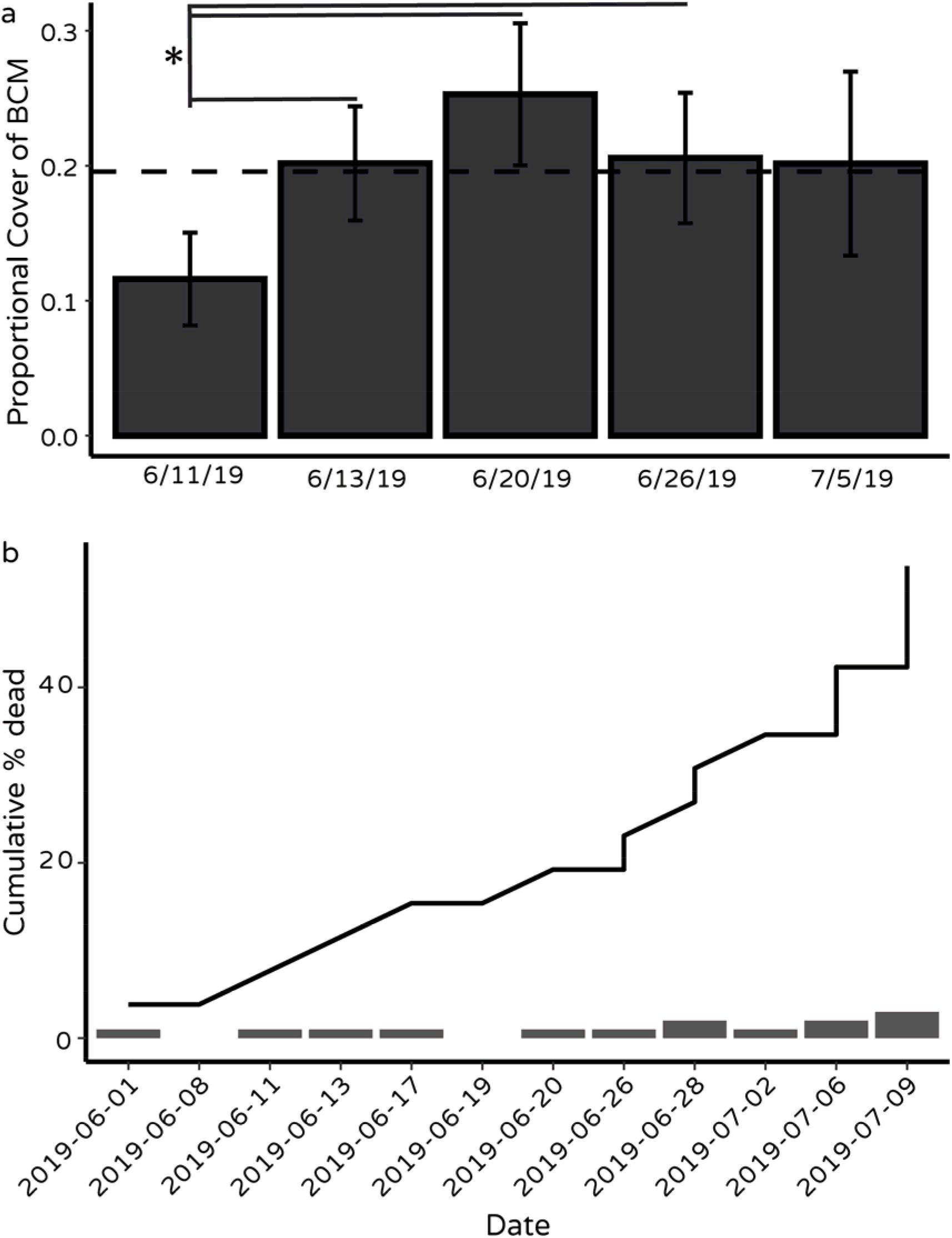
a, Barplots of site-scale proportional cover of cyanobacterial mats across sampling dates. Error bars correspond to standard error around the mean for each sampling date. Horizontal dashed line denotes mean cover across sampling dates. b, Line plot of cumulative percent of component communities dead across sampling dates. Rug barplots show individual mat death counts across dates (scaling axis not shown).

Alongside documented metacommunity stability across the study period, component communities showed remarkable volatility (Fig. 1b). At the end of the 49-day study period, 53.8% of the tracked cyanobacterial mat communities (considering only unmanipulated mats; 14 / 26 mat individuals) had died across 10 unique death dates (Fig. 1b), indicating a high degree of asynchrony in the dynamics of individual component communities comprising this mat metacommunity.

### Experimental wounding does not increase likelihood of mat death, and decreases predicted instantaneous mortality rate of mat communities

To test for an effect of macroscale disturbance (most closely approximating direct effects from predation by consumptive reef fishes) on the dynamics of benthic cyanobacterial mats, 27 mats were experimentally cored in the field, representing a ‘wounding’ treatment over unmanipulated mats. Motivated by previous work demonstrating a strong antagonistic effect of predation pressure on the establishment and persistence of experimental microbial mats (Fenchel 1998), we hypothesized that enhanced predation would result in decreased reef cyanobacterial mat persistence via a strong community response to top-down pressure. We found little support for this hypothesis, with no significant difference in the overall probability of mat community death over unmanipulated mats (GLM, F = 0.114, df = 1, p = 0.74; Fig. 2a). Interestingly, however, cyanobacterial mats in the coring treatment were predicted to survive longer, having a significantly lower instantaneous mortality rate (−0.012 [-0.024 - -0.005 97.5% CI]) than mats that were left unmanipulated (−0.023 [-0.041 - -0.011 97.5% CI]; GLM, dev = 8, df = 2, p < 2.2e-16), suggesting that moderately increased predation pressure increases mat longevity (Fig. 2b).

**Figure 2.**
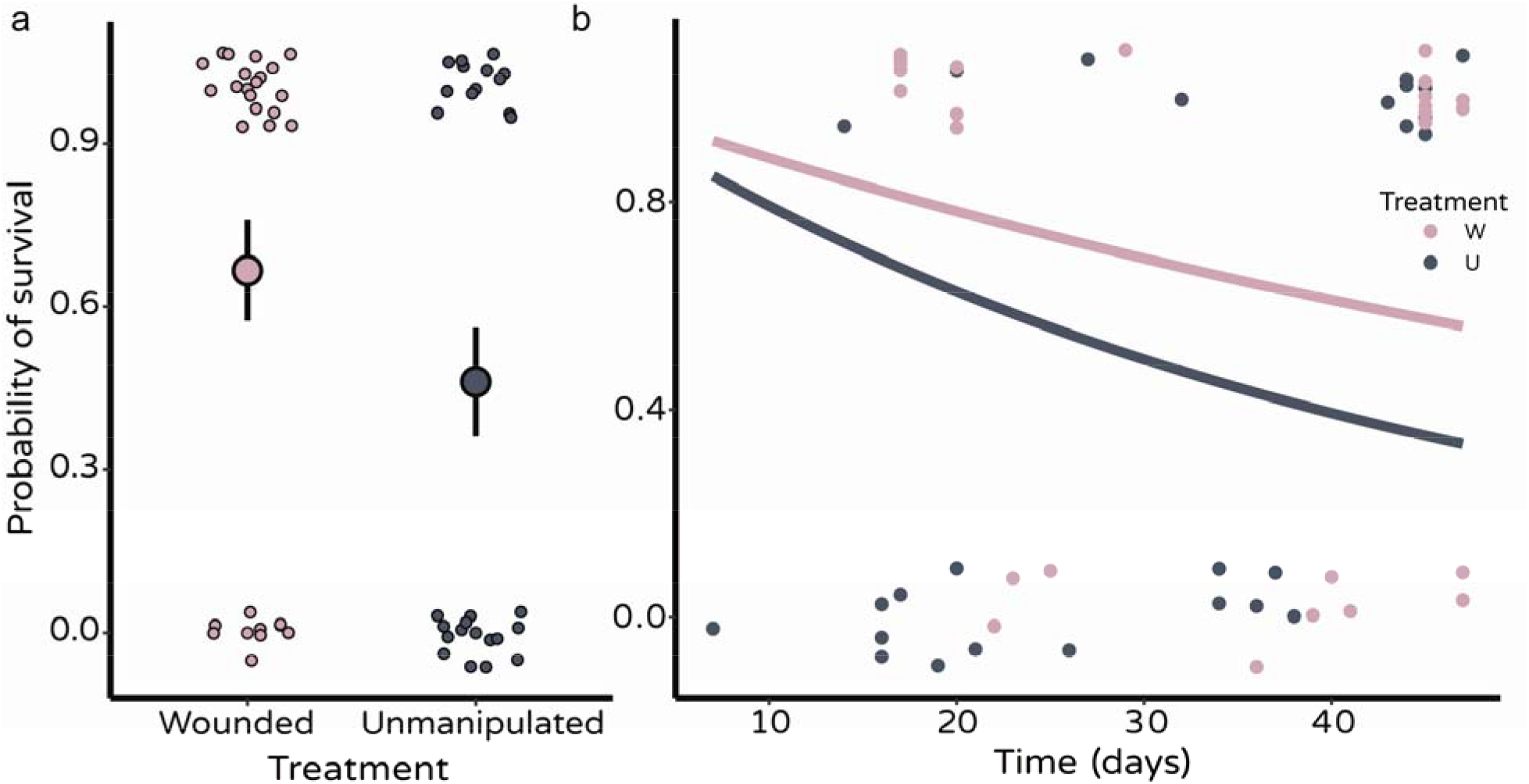
a, Probability of individual community death by treatment inclusion. Large circles denote mean probability of a mat community dying in each treatment; error bars represent SE. Individual points are binomial outcomes corresponding to if a mat died during the study period (0) or not (1). b, Exponential survival curves from benthic cyanobacterial mats subject to natural grazing pressure (Unmanipulated [U]; gray) vs mats experimentally wounded (Wounded [W]; pink) demonstrating reduction in instantaneous mortality rate with increased predation. Raw points (jittered vertically) show individual mat outcomes (binomial; 1=survival, 0=death).

### Macropredation pressure is heterogenous among mat communities, and varies by predator identity

Diver-independent fixed-point behavioral assays were deployed on 22 spatially distinct cyanobacterial mat communities across five different reef sites to identify relevant sources of macropredation and quantify rates of predation pressure. A total of 11 different fish species were documented to have foraging behavior that impacted cyanobacterial mat biomass, with both initial phase and terminal phase individuals from 3 species observed grazing on cyanobacterial mat biomass (Fig. 3). While bite pressure differed significantly by fish species identity (chisq = 39.2; df = 13; p = 0.0002), no significant difference among sampled sites was detected (chisq = 21.59; df = 4, p = 0.81; Figure S5). Predation pressure among mats, however, was highly variable (72.8 mean bites per mat ± 111.2 bites SD; GLMM random effect variance 3.12 ± 1.77 SD), overall suggesting heterogenous macropredation pressure from reef fishes within (i.e. heterogenous among individual mat communities), but not necessarily among, mat metacommunities (i.e. relatively homogenous among sites). Mean bite rate (bites*min^-1^) across all species taken on cyanobacterial mat communities was 0.23 ± 0.92SD. Though greatest in frequency (Fig. 3), bites taken by the Bicolor Damselfish (*Stegastes partitus*) were of lesser magnitude (i.e. did not remove as much visible mat biomass) than bites by other fish species.

**Figure 3.**
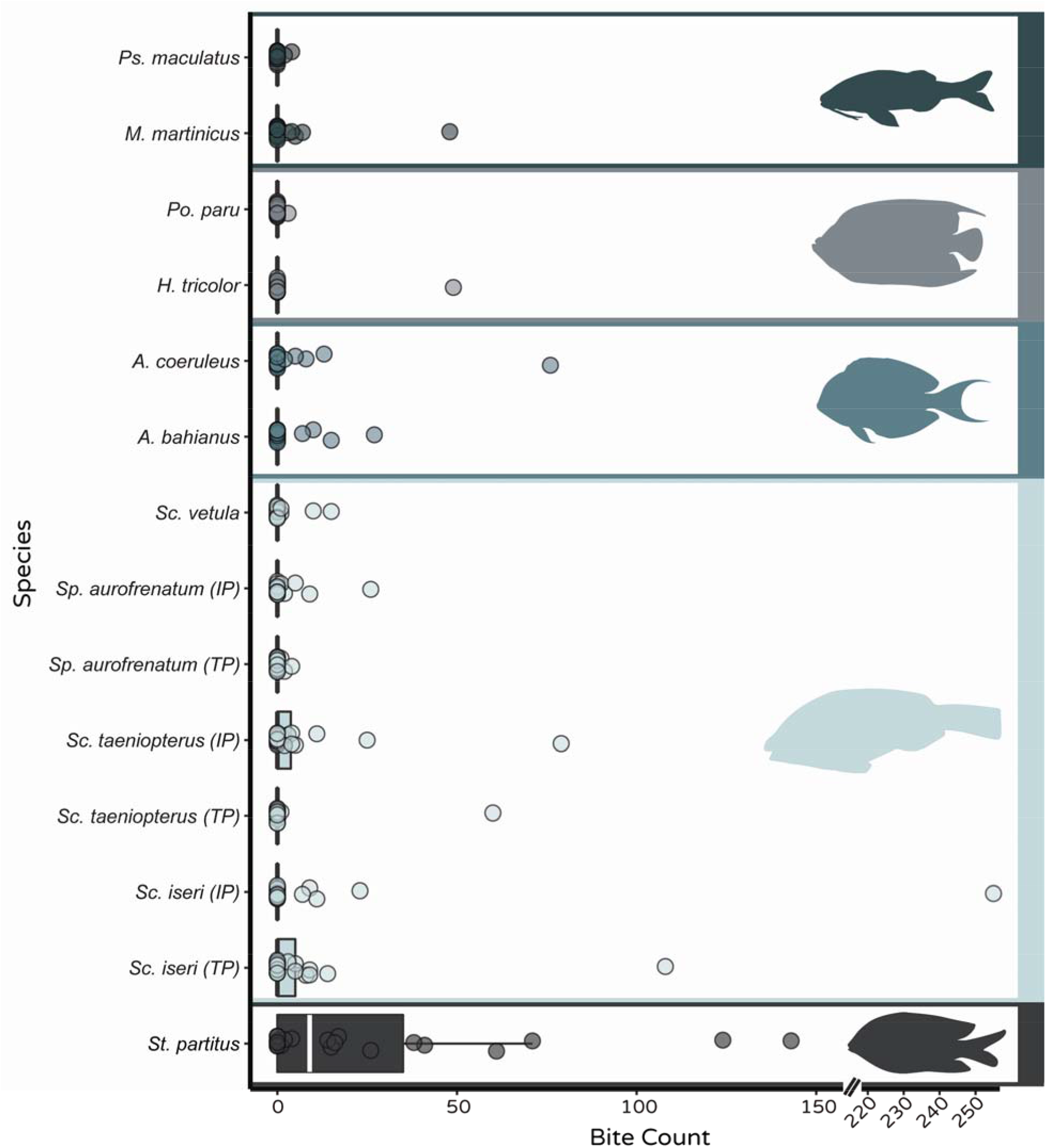
Boxplots of bite count (#) by fish species. Data for *Pseudupeneus maculatus* and *Mulloidichthys martinicus* represent total number of forays that disturbed cyanobacterial mat biomass and not bites (see Section 5.2. Materials & Methods). *TP* and *IP* designation next to species names on y-axis denote *Terminal Phase* and *Initial Phase* respectively. Colors and silhouettes broadly group species by taxonomic family (i.e. Mullidae, Pomacanthidae, Acanthuridae, Scaridae, Pomacentridae from top to bottom). Fish silhouettes are from *R::fishualize*.

Similarly, bites taken by larger predators such as the French Angelfish (*Pomacanthus paru*) and the Queen Parrotfish (*Scarus vetula*), though fewer in frequency, were of much greater magnitude and removed significant biomass with each bite. The number of bites taken by initial phase Striped Parrotfish (*Scarus iseri*; mean 13.8), Princess Parrotfish (*Scarus taeniopterus*; mean 6.2) and Redband Parrotfish (*Sparisoma aurofrenatum*; mean 2.0), were generally higher than those taken by terminal phase individuals (means 7.3, 2.8, and 0.3, respectively). However, again, an effect of species changes the magnitude of each bite, with greater biomass removal from terminal phase individuals.

### Moderate dispersal conveys rescue effect to mat metacommunities even with strong top-down control

A metacommunity cellular automaton model was employed to explore how different scenarios of bottom-up forcing, top-down control, and dispersal drive mat metacommunity dynamics. Scenarios at naturally unlikely extremes (i.e. zero top-down forcing or immense bottom-up control [zero growth]) showed outcomes well aligned with *a priori* assumptions, either saturating the metacommunity at 100% benthic cover or ending with the metacommunity going extinct, persisting at 0% benthic cover (Fig. 4). Model outputs suggest that even moderate levels of top-down control from predation can significantly reduce the relative benthic cover of mat metacommunities, with high levels of top-down control almost always driving metacommunity extinction without dispersal (Fig. 4). Interestingly however, strong dispersal potential conveys a rescuing effect on metacommunity persistence even in scenarios with relatively strong top-down and bottom-up control (mean equilibrium proportional abundance ∼1.6% under High Predation, Low Growth, High Dispersal, and ∼1.2% under High Predation, Zero Growth, and High Dispersal; Fig. 4). Moderate dispersal can still promote persistence, but at extremely low cover under strong control scenarios (∼0.16% cover under H_Z_M scenario). Further, under more realistic scenarios (i.e. Moderate top-down and bottom-up control), strong dispersal potential can maintain metacommunity cover at values close to the starting abundance, mirroring patterns empirically observed in the field during this study period (Fig. 1, 4). This suggests that strong dispersal is an important mechanism driving mat metacommunity dynamics on reefs.

**Figure 4.**
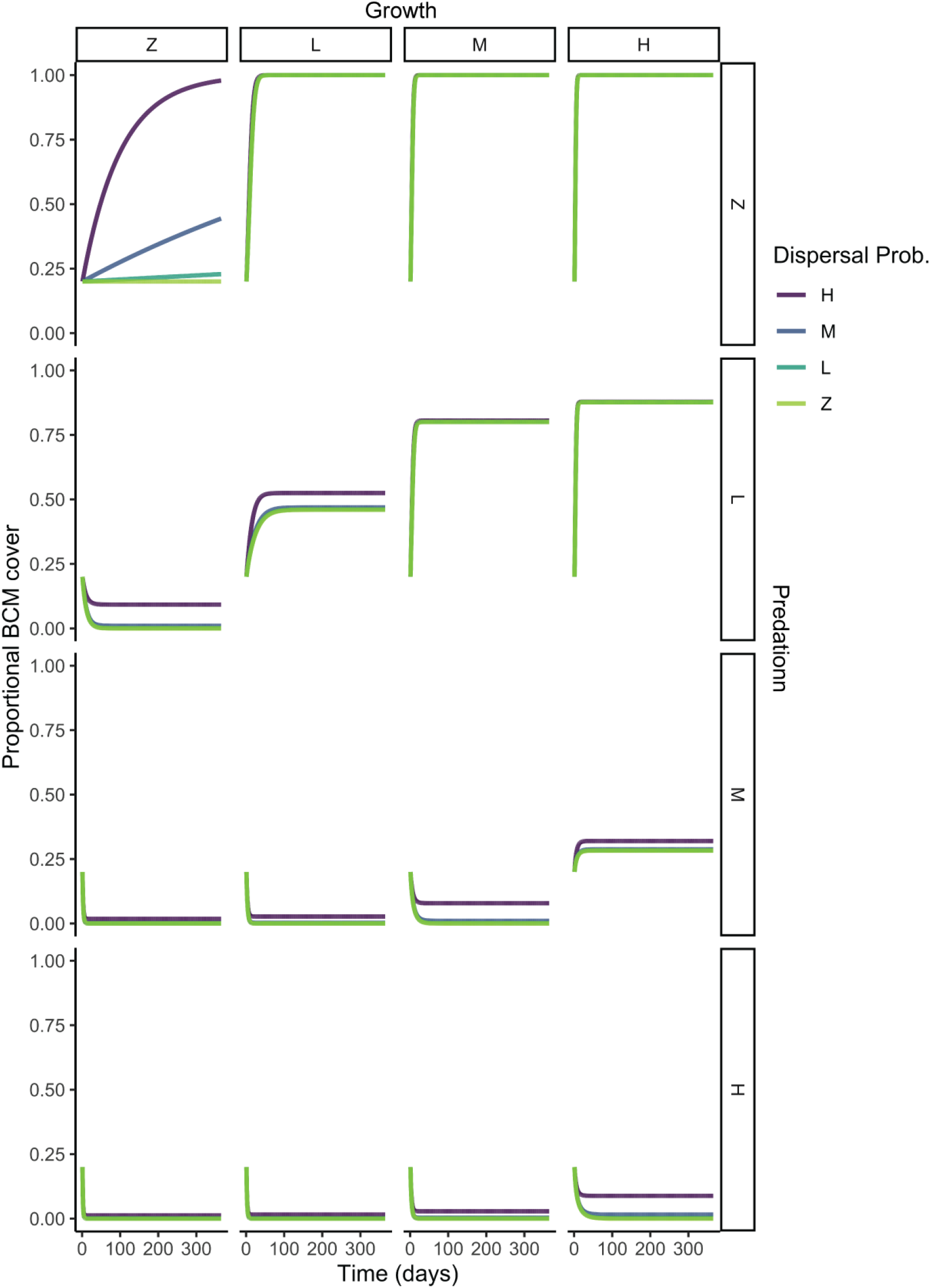
Outputs from model simulations showing mean (from n=100 separate model simulations each) trajectory of cyanobacterial mat cover across the 1.25km^2^ model grid area over 365 days. Plots are faceted by different parameterizations of growth rate across the X axis, and different parameterizations of predation rate across the Y axis. Different parameterizations of dispersal rate are encoded in different line colors. Lines do not represent statistical regressions, and are present to aid in trend visualization.

## DISCUSSION

### Decoupling of local and regional scale patterns

In comparison to persistent mat metacommunity stability, component mat communities displayed remarkable instability in physical space (Fig. 1). However, the temporal pattern of senescence of individual communities was highly asynchronous, occurring across 10 unique death dates (Fig. 1b). This asynchrony can strongly decouple patterns in community and metacommunity dynamics and mitigates the correlative strength between local community extinction risk and regional metacommunity extinction risk. Asynchrony in substructure component dynamics (i.e. decorrelation of population and community extinction and recolonization across axes of space and time) has long been appreciated as an important systemic stabilizing force in both metapopulation and metacommunity ecology, promoting stability in metacommunity biomass and ecosystem function by decreasing intra-scale variability of abundances (Paine and Fenchel 1994, Holyoak and Lawler 1996, Wilcox et al. 2017). Predation pressure can be an important force toward preventing synchrony among communities and promoting compensatory oscillatory dynamics in metacommunities, even in systems with strongly synchronizing dispersal (Howeth and Leibold 2013). However, the extent of these effects depends strongly upon predator niche, distribution of predation pressure in time and space, and context of abiotic environmental variability, as predators can antithetically promote species and spatial patch synchrony via spatial coupling of dynamics (Howeth and Leibold 2010, 2013, Firkowski et al. 2022).

The documented spatial heterogeneity in experienced predation pressure from generalist mobile reef fishes across mat communities may significantly contribute to the maintenance of asynchrony in the dynamics of component communities in this cyanobacterial mat system via the creation of spatiotemporal complementarity in patterns of mat senescence (Fig. 1b). The need to further understand and incorporate trophic interactions into our understanding of mat metacommunity dynamics is critical, especially considering the increasingly dramatic environmental fluctuations experienced on reefs (i.e. massive pulse nutrient loadings; Firkowski et al. 2022). Trophic interactions may be a strong local-scale generator of asynchrony in community dynamics despite regional spatially synchronizing effects resulting from spatially correlated responses to environmental fluctuations driven by relatively low beta diversity among local mat communities (Cissell and McCoy *unpublished manuscript*), helping to maintain persistent mat cover at the scale of reef site. These data provide critical context toward explicitly incorporating trophic interaction strengths and variability into predictions of site-level and regional dynamics (Paine 1992, 2002).

### The role of disturbance and the potential for top-down control

Disturbance events from both physical and biological processes are critical driving forces in the creation of spatiotemporal variability in the dynamics of natural communities (Sousa 1984). Indeed, local-scale interactions, including predator-prey linkages (Livingston et al. 2017), are known to scale to the structure and function of entire communities and metacommunities (Baiser et al. 2013). Predatory interactions seem to be relevant mechanisms of disturbance for driving patterns across scales of organization in benthic cyanobacterial mats on coral reefs, from intra-mat scales (i.e. population-specific interactions from viral specialist predators (Cissell and McCoy 2022a), to whole community scales (i.e. generalist grazing from reef fishes [Fig. 3; Cissell et al. 2019]). Though strongly dependent upon the structure of the competitive network (Paine 1971), specialist predation is generally thought to have smaller direct effects on community and metacommunity stability (Howeth and Leibold 2010), which might suggest that predation from reef fishes on mats may play an outsized role over viral predation in driving empirically observed community instability. We documented a diverse suite of grazers contributing to total mat community macropredatory pressure (Fig. 3). Because of differences in gape size and grazer physiology, significant differences in total foraging on mats among these different species of reef fishes suggests mats experience variability in both predation frequency and magnitude, which may in turn result in divergent physiological impacts to the mat intrinsically coupled to grazer identity (i.e., consumer trait-mediated effects).

Microbial mats that form in other ecosystems are generally long-lived owing to their capabilities for nutrient recycling and retention (e.g., months - years; Doemel and Brock 1977, Bolhuis et al. 2014, Stal et al. 2019). Reef cyanobacterial mats possess similar mechanisms promoting nutrient retention within the mat system (Cissell and McCoy 2021; Cissell & McCoy *unpublished manuscript*), suggesting the feasibility of an *a priori* hypothesis predicting individual mat persistence on reefs and a minimization of the predominance of bottom-up control for mature mat community decay. This, in-turn, suggests that observed patterns from dynamism in mat abundance could be attributed to top-down forcing. This is not to say that bottom-up forcing does not influence mat growth dynamics, as bottom-up processes have long been implicated in mediating cyanobacterial mat growth rates (Kuffner and Paul 2001), but rather suggests that mat community dissolution and site-wide abundance of distinct mat individuals can respond to top-down forcing. The striking instability of mats that we have observed empirically over this relatively short time-period suggests that mat communities readily respond to top-down disturbance from predation. This might imply that natural levels of predation experienced by mat communities impart strongly unidirectional outcomes on controlling mat communities, which is supported by the presence of a strong effect of top-down control in driving reductions of mat metacommunities in model simulations (Fig. 4).

The surprising outcomes from the coring experiment complicate the generalization of the role of top-down forcing in driving mat community dynamics by suggesting non-linearity in the response of mat prey to predation pressure. These experimental data, though, may help to contextualize the relative magnitude of naturally occurring predation pressure. The nonequilibrium maintenance of biodiversity has long been thought to be dependent upon the frequency and magnitude of disturbance acting upon the system (Connell 1978). Indeed non-specific disturbance (including from generalist predators), in addition to density-dependent specialist predation (Paine 1971, Hewson et al. 2003, Thingstad et al. 2014), can be critical in the promotion of species coexistence by preventing competitive exclusion (Sousa 1979). Cyanobacterial mats are mutualistic consortia that depend upon metabolic coupling among physiologically and trophically distinct populations for community persistence (Cissell & McCoy *unpublished manuscript*). Maintenance of biodiversity, then, is critical for mat persistence, which would suggest some intermediate level of disturbance may benefit the longevity of individual mat communities.

Indeed, maintenance of high intra-mat biodiversity likely stabilizes mean community productivity and buffers variability in critical community functions in the face of relatively stochastic but extreme-in-magnitude environmental fluctuations from anthropogenically-derived terrestrial-based nutrient subsidies (i.e. the insurance hypothesis; Yachi and Loreau 1999, Ford et al. 2017). Stabilizing interactions (both immediate and cascading from pulse restructuring of interaction networks [Fazzino et al. 2019]) from exogenous predation may further benefit mat longevity by buffering mat communities against the destabilizing effects of extensive cooperation (May 1972, Allesina and Tang 2012, Coyte et al. 2015). The significantly increased longevity of those mats in the experimental coring treatment qualitatively supports this biodiversity maintenance hypothesis (Fig. 2b) and suggests that natural predation regimes experienced by mats on the reefs in Bonaire may be on the low side of intermediate. This may at first appear counterintuitive to the outputs from the model simulations, which suggest that moderate predation pressure (including empirically informed levels of fish predation) imparts significant mortality on mats and depreciation of metacommunity abundance, which would imply decreased longevity of component communities. However, importantly, these model simulations necessarily exclude the existence of interactive effects among trophic events and growth rate (independently estimated and not dynamic) and were limited in assuming homogeneity of predation risk across axes of time and space. This homogeneity applied even for high-in-magnitude low-in-frequency disturbance events arising from the foraging behavior of goatfishes. Indeed, the ‘Low’ predation model simulations imposed control over mat metacommunity abundance from saturating at 100%. In other words, the ‘Medium’ predation scenario in the model assumed homogeneity of pressure across both time and space, which may have pushed these modeled scenarios beyond what may be typically viewed as ‘intermediate’ levels of disturbance. Additionally, model outputs approximated cover estimates at the level of metacommunity and did not track the dynamics of individual mat communities. In this way, the model more closely approximated the resolution of typical monitoring surveys and cannot make inference on the ability of the different parameterizations of predation risk to entirely remove individual mat communities.

Generalized metabolic cooperation may also be predicted to be detrimental to community stability by imposing an interdependence of population densities (May 1972, Allesina and Tang 2012, Coyte et al. 2015, Hoek et al. 2016). If top-down control disrupts cooperative interactions, especially asynchronously (Paine and Fenchel 1994), then metabolic cooperation may become a double-edged sword due to the repercussions of decoupled cooperative interactions. Macropredation pressure on mats is necessarily heterogenous across an individual mat community ‘landscape’, and we found evidence to support that micropredation from viruses, too, is heterogenous across a mat community. Predation, then, could change local-scale linkages within mat communities, as is observed in experimental microbial metacommunities (Livingston et al. 2017) and synthetic metabolically-coupled microbial networks (Fazzino et al. 2019), interacting with niche-based processes to drive metabolic asynchrony and promoting overall community instability. Instances of mat senescence have been observed radiating out from apparent bite scars, suggesting the potential for dynamics to be independent of discrete trophic events and instead linked to emergent higher order effects from cascading trophic influence (Figure S6). Within-mat patch dynamics could arise when space is opened from macropredation for recolonization (local-scale disturbance), which may interact with heterogeneity in predation frequency to promote intra-mat landscape mosaics of taxonomic and functional richness (Sousa 1979, Paine and Levin 1981; Figure S7). Similar patchy patterns of diversity have previously been demonstrated in hot-spring cyanobacterial mats during recolonization following experimental disturbance (Ferris et al. 1996, 1997). Primary recolonization of bare substrate is likely predominantly pioneered by cyanobacterial species (Stal et al. 1985), likely shifting total mat stoichiometry in C:N:S via an apparent cyanobacterial bias.

Niche plasticity within cyanobacterial mat component populations, however, may dampen any negative effects of stoichiometric shifts (Cissell & McCoy *unpublished manuscript*). Linkages among metabolic asynchrony, biodiversity maintenance, predation pressure, and community spatiotemporal asynchrony warrant further exploration across diverse systems with differing levels of predation pressure (i.e. varying biotic and abiotic context; Cissell et al. 2019, Ford et al. 2021, Ribeiro et al. 2022) to better understand the influence of predation in driving the structure and dynamics of cyanobacterial mat communities and metacommunities.

### The rescuing role of dispersal

Our simulations predict that strong dispersal can meaningfully rescue mat metacommunities from extinction even under scenarios of strong top-down pressure and growth-limiting bottom-up control (Fig. 4). Dispersal may be especially crucial in supporting mat metacommunity persistence during episodic periods of relatively stable top-down pressure and strong bottom-up control in generally oligotrophic reef environments experiencing periodic land-based input (Brocke et al. 2015a, den Haan et al. 2016, Ford et al. 2017). The results of our theoretical model contribute to explaining the apparent decoupling of mat cover from targeted management actions tailored to increasing bottom-up control (i.e., limiting nutrient inputs) periodically observed (e.g., high documented high mat cover despite low nutrient loads and low Anthropogenic influence; Brocke et al. 2015a), as our results suggest that strong dispersal would facilitate this persistence at the metacommunity scale during periods of strong control on individual community persistence imposed by management practices (Fig. 4). Dispersal linkages were previously suggested to be present among cyanobacterial mat communities from molecular evidence demonstrating strong genomic conservation among spatially distinct mats (Cissell & McCoy *unpublished manuscript*). For cyanobacterial mats, predation, especially from mobile predators, may be an important mechanism promoting mat dispersal (Cissell et al. 2019, 2022). Predation by reef fishes has previously been suggested as an important vector of dispersal for endosymbiotic dinoflagellates of corals via grazing and subsequent fecal deposition (Grupstra et al. 2021), and consumer-mediated dispersal is more generally thought to be an important, yet understudied component of microbial community assembly and dynamics (Grupstra et al. 2022). In mats, foraging by reef fishes, and especially goatfishes, may also mechanically dislodge mats from the sediment surface into the water column to be dispersed via water movement, in addition to potential dispersal via fecal deposition. Further work is needed to better understand mechanisms of cyanobacterial mat dispersal across reef landscapes, as well as the temporal scope of these rescuing effects (duration of strong control and maintenance of cover from dispersal linkages).

Taken all together, mats living at sediment surface may experience interesting and unique trade-offs. One side of the tradeoff could be likened to a pseudo-reversal of the paradox of enrichment (Rosenzweig 1971) – treating surface sediment as the enriched prey resource, then prey capitalization on increased resource availability (i.e. growing primarily at the surface) may directly lead to prey population collapse (reversal) from density-dependent predator response. In this sense, minimizing surface-sediment biomass available for opportunistic or targeting grazing would increase individual mat longevity. However, if predation is critical for mat dispersal, component community instability from predation may be necessary for long-term metacommunity persistence – suggesting mat dynamics may be more closely akin to the classic ‘blinking lights’ models of metapopulation ecology describing the dynamics of short-lived populations (Levins 1969).

### Top-down vs. bottom-up control and the importance of scale

The focus on documenting the relative influence of top-down forcing on mat dynamics herein is not to ignore or minimize the empirically documented importance of bottom-up factors (primarily sediment-bound nutrient context), and physical factors more generally, in governing the dynamics of cyanobacterial mats on coral reefs (Kuffner and Paul 2001, Albert et al. 2005, Hallock 2005, Ahern et al. 2007, Brocke et al. 2015b, Tebbett et al. 2022). Indeed, many of the documented responses to top-down forcing likely interact with other highly relevant physical driving factors such as light availability, temperature, hydrodynamics, and sediment/intra-mat redox conditions. Regional-scale disturbance from physical wave energy and general hydrodynamic turbulence on sediment topography (including scouring) have previously been suggested as important determinants of cyanobacterial mat distribution on coral reefs (Thacker and Paul 2001, Tebbett et al. 2022). The formation of sand ripples from water motion may altogether preclude mat formation, and disrupt patterns of persistence in mature mats, contributing to site-scale patterns in mat distribution and persistence. The opening of channels from local-scale top-down trophic events within the generally cohesive structural matrix binding mat communities may create areas more susceptible to physical disruption from water movement (from the formation of non-cohesive edges), which may interactively contribute to mat dynamism. Such trophic-mediated intra-mat channels may further alter redox conditions in the underlying sediment, as cover of reef cyanobacterial mats (Brocke et al. 2015a), and macrophyte cover more generally (Boros et al. 2011), are shown to have strong linkages with sediment redox potential. Shifting redox potential may strongly change the habit of physical mat manifestations and may drive mat communities deeper into the underlying sediment or promote the full disintegration of conspicuous mat structure from shifting nutrient settings.

Though light availability at the sediment surface would maximize cyanobacterial photosynthetic efficiency, living subsurface may confer some benefits as described above in addition to protection from exposure from ultraviolet radiation which may damage critical photosynthetic machinery (Garcia-Pichel and Bebout 1996). Further, mat size at the surface across its lifespan is likely an emergent result of interactions among predation pressure, hydrodynamic setting, and benthic nutrient supply, with a likely positive covariance among areal extension and experienced predation pressure (primarily from reef fishes).

This discussion raises an additional interesting point concerning the manifestation of mat ‘death’ presented herein. Our assessment of mat death was reliant upon conspicuous manifestations that were observable to the naked eye, namely the disappearance of conspicuous mat matrix from the surface sediment. Indeed, viable microbial cells may still be present from the mat community but may not be visible or may have migrated subsurface. Surface vs. subsurface manifestations of mats may be likened to ‘life-stages’ of mats that are coupled to immediate abiotic and biotic context and history. Further exploration of cell integrity / viability associated with the disappearance of conspicuous surface level cyanobacterial mat biomass is necessary to resolve how closely the disappearance of conspicuous mat matrix can be associated with true ‘death’ of individual mat communities.

Collectively, these data indicate that both top-down and bottom-up forcing should be considered when constructing general frameworks toward understanding the dynamics of cyanobacterial mats on coral reefs, and that scale - both in ecological organization and physical space - likely matters when determining the relevance of trophic forcing directionality. Understanding how trophic dynamics interact with mat cover across scales is critical toward the creation of well-informed management strategies for controlling standing mat biomass, and for preventing the formation of new mat biomass on reefs (Ethan C. Cissell and McCoy 2022).

### Synthesis and future recommendations

Here, we showed that coral reef benthic cyanobacterial mat metacommunities are temporally stable despite dramatic instability in component community patches across short time scales (Fig. 1). This decoupling of extinction risk across scales is likely linked to spatiotemporal asynchrony in the dynamics of component patches, which may, in part, be driven by heterogeneity in predation pressure across communities (Fig. 3). Predation pressure may not drive unidirectional outcomes in benthic cyanobacterial mat dynamics, though, with field experiments demonstrating a decreased instantaneous mortality rate of mats in response to increased predation pressure (Fig. 2b), potentially mediated via disturbance-diversity relationships. Dispersal may also play a strong role in driving the dynamics of mat metacommunities and is predicted in simulation modeling to confer a rescue effect to mat cover under scenarios of strong top-down and bottom-up control (Fig. 4). These data establish critical baselines and generate hypotheses relevant to ecologists and managers alike on the processes maintaining cyanobacterial dominance of coral reefs. We recommend that future work on cyanobacterial mat metacommunities focus increasingly on 1) explicitly understanding spatiotemporal variability in fluctuations among taxonomically (and, perhaps more interestingly, functionally) distinct populations within local communities and among similar populations within the broader metapopulation toward further parsing the implicit hierarchical variance structure across these distinct yet linked levels of ecological organization (*sensu* Wang and Loreau 2014, Hammond et al. 2020); 2) identifying the relevant mechanisms and spatial structure of dispersal (as previously recommended in Cissell et al. 2022); 3) working toward understanding the link between the stability in metacommunity biomass reported here, and both the mean and spatiotemporal variance in mat metacommunity functional ecology (i.e., metacommunity scale DOC release, N_2_ fixation, etc.; Brocke et al. 2015b, 2018, Cissell & McCoy *unpublished manuscript*); and 4) working toward quantifying the linkage strengths of trophic interrelations reported herein using classic removal experiments to further disentangle the importance of coupled dynamical trophic modules vs competitive linkages previously reported (Thacker et al. 2001, Puyana et al. 2019) in structuring benthic cyanobacterial mat metacommunity demography on coral reefs (Paine 1980).

## ACKNOWLEDGMENTS

Funding for this work was provided by a gift from the Tatelbaum Fund and start-up funding from Florida State University to S.J. McCoy. E.C. Cissell was supported by a National Science Foundation Graduate Research Fellowship (Grant Number: 074012-520-044116) and a Mote Research Assistantship from the William R. and Lenore Mote Eminent Scholar in Marine Biology Endowment at Florida State University. We thank R.L. Francisca and C.E. Eckrich at Stichting Nationale Parken (STINAPA) Bonaire and Rijkswaterstaat Ministerie van Infrastructuur en Waterstaat for research permissions for conducting this work within the Bonaire National Marine Park. We extend our thanks to J.C. Manning, I. Basden, M. Dziewit, and B. Clark for assistance in the field, and L. Kury in the analysis of benthic photoquadrat data. We thank C. Peters for supporting the diving component of this work. We thank D.K. Okamoto for help with statistical analyses. Finally, we thank J.R. Cissell, K.M. Majzner, J.C. Manning, K.L. Dobson M. Huettel, D.K. Okamoto, K.M. Jones, and D.R. Rokyta for discussions and feedback on earlier versions of this manuscript.

## AUTHOR CONTRIBUTIONS

E.C. Cissell conceived of and designed the study with significant contributions from S.J. McCoy. E.C. Cissell led all data collection in the field, and all in-situ data analyses. S.J. McCoy significantly contributed to in-situ data analysis. E.C. Cissell drafted the original manuscript with significant contributions from S.J. McCoy. Both authors contributed significantly to manuscript revision.

## CONFLICT OF INTEREST STATEMENT

The authors declare they have no known competing financial interests or personal relationships that did or could have appeared to influence the work reported in this paper.

## Notes

### Competing Interest Statement

The authors have declared no competing interest.

### Summary of Updates

This version of the manuscript has been revised following Editorial Board comments

