## Supporting Information PDF for "Predation, community asynchrony, and metacommunity stability in cyanobacterial mats"

**Supplementary Information Contains:**

**___**

Supplementary Discussion,

**16** Supporting Figures,

&

**1** Supporting Tables

___

**SUPPLEMENTARY DISCUSSION**

We join recent calls in advocating for the importance of understanding and documenting system-specific natural history in addition to applying generalizable theory toward holistically asking and contextualizing hypothesis-driven research (Travis 2020). Toward that end, what follows is a curated selection of relevant natural history observations made during our study period that further contextualize the patterns and processes presented herein, and which hopefully will generate and inform future hypotheses about relevant pattern-generating mechanisms in the dynamics of benthic cyanobacterial mat metacommunities.

In Bonaire, the foraging behavior of the goatfishes *M. martinicus* and *Ps. maculatus* can both directly and indirectly disturb benthic cyanobacterial mats. As a direct result of foraging in sediment underlying mats, *M. martinicus* individuals can disrupt mat structure, and remove sections of the mat from the benthos. This disturbance to mat biomass results primarily from the ‘shovel’ and ‘push’ feeding modes, however ‘skimming’ behavior also disrupts mat structure. These individuals are likely not foraging on the mat itself, as goatfish primarily forage on benthic invertebrates (Krajewski et al. 2006). Goatfish targeting of sediment underlying mats for forays may be promoted by the forced vertical migration of sediment infauna toward the sediment surface from mat-promoted sediment anoxia (Brocke et al. 2015), as previously suggested by Cissell et al. (2019) when discussing the targeting of mats by other mobile omnivorous reef fishes (primarily *Po. paru* [French Angelfish]). In addition to the direct effects of foraging on sediment underlying mats, we observed that *M. martinicus* individuals can partially or completely bury a mat in sediment by foraging nearby, as we directly observed (Figure S8). Indirectly, this rapid sedimentation of mat surface area should negatively impact mat photosynthetic yield and efficiency – the dominant pathway of primary carbon acquisition in cyanobacterial mats (Cissell and McCoy 2021) – as is observed in other photosynthetic taxa (Izagirre et al. 2009). This would necessitate relatively rapid vertical migration of the mat structure to overcome the rapid sedimentation event, or rapid shifts in the predominance of enzymatic machinery, and could even disrupt carbon balances within mats.

On numerous occasions during our observational period, we observed *Elysia crispata* (Lettuce Sea Slug) individuals on benthic cyanobacterial mats overgrowing both hard substrate and sediment (Figure S9). Although in this study we did not directly quantify consumption of mats by *E. crispata* individuals and so we cannot say how circumstantial these associations were, multiple lines of evidence suggest their ability and potential to consume benthic cyanobacterial mats. *E. crispata* are members of the Sacoglossa, marine sea slugs within the Opisthobranchia, and are specialist herbivores that feed on marine algae and seagrass (Clark and Busacca 1978). Many species within this group retain functional, diet-derived plastids in a process called kleptoplasty. This unique phenomenon is restricted in metazoans to members of this gastropod taxon. Among the Sacoglossa, the genus *Elysia* shows the most diverse range of organisms targeted, with *E. crispata* targeting a notably wide range of food organisms (Händeler and Wägele 2007). Other Opisthobranchs (*Stylocheilus* spp.) are known specialist consumers of benthic cyanobacterial mats on both Caribbean (Capper and Paul 2008; Arthur et al. 2009) and Pacific coral reefs (Nagle et al. 1998). Furthermore, since *E. crispata* lack shells, benthic cyanobacterial mats may be a rich source of toxic secondary metabolites that these individuals could sequester and use to decrease their susceptibility to predation, which has been previously observed in Sacoglossans with noxious secondary metabolites derived from siphonalean green algae (Hay et al. 1989). Some Sacoglossans show ontogenetic shifts in dietary targets (Curtis et al. 2006), and therefore consumption of benthic cyanobacterial mats by *E. crispata* individuals may vary across its development. We recommend that future work be directed toward understanding the interactions between *E. crispata* and benthic cyanobacterial mats, including kleptoplasty, the assimilation of toxic secondary metabolites from the mat, and the level of disturbance the mat may sustain resulting from these potential interactions.

*St. partitus* (bicolor damselfish) individuals were found associated with 19 of the 22 cyanobacterial mats observed for fish foraging observations (~86%). During the observation periods, we observed numerous antagonistic chases against heterospecific grazers that resulted in predator deterrence from grazing with associational retention of the damselfish individual(s) near (often directly above) the conspicuous mat matrix throughout the entire observation period, suggesting these cyanobacterial mats fall within the territories of these *St. partitus* individuals (Myrberg 1972). These territories were not exclusive from conspecifics, however, with a maximum of three *St. partitus* individuals being observed simultaneously within frame over a single observed mat community (mean 1.8 individuals per mat community). Bites on mat communities by *St. partitus* individuals were quite frequent (Fig. 3) but were small in magnitude. Other damselfishes, including other species within the genus *Stegastes*, are well known for their ‘farming’ behavior with algal turfs – a mutualism that promotes algal growth via nutrient subsidies and protection from predators via behaviorally-mediated exclusion while the farming damselfish receives a consistent nutrient-dense food source (Hata et al. 2010; Blanchette et al. 2019). We hypothesize a similar relationship with emergent cyanobacterial mats falling within the territories of bicolor damselfish, which may contribute to the documented heterogeneity in predation pressure from reef fishes across component communities within a mat metacommunity, suggesting that the biotic context of cyanobacterial mats is likely a critical contributor to variability among individual mat community dynamics. Maintaining lawns consisting largely of benthic cyanobacterial mat cover may help increase nitrogen intake rates of *St. partitus* individuals given the relatively low C:N ratios of some mat-forming marine benthic cyanobacterial species relative to the upright filamentous algal species that are often farmed by damselfishes (Atkinson and Smith 1983; Hata and Kato 2002). Alternatively, the bites we observed during our video observations may represent individuals attempting to weed out and promote exclusion of these cyanobacterial mats from their territory in an attempt to maintain cover of non-cyanobacterial mat algal taxa (Hata and Kato 2004). Further work should attempt to differentiate among these competing behavioral hypotheses to better understand the relationship between farming damselfish and benthic cyanobacterial mats.

The immediate physical context of reef structure surrounding spatially distinct mat communities may play an outsized role in determining experienced predation pressure and driving variation in predation among mat communities. Structural complexity has long been thought to mediate the relative effects of competition vs. predation in driving community diversity (Menge and Sutherland 1976) by mediating predation intensity and interacting with foraging efficiency, preventing overexploitation (Paine 1974). These same arguments logically extend to cyanobacterial mat communities from the perspective of both community diversity and biomass (links discussed in more detail in main manuscript discussion). The physical context of mats also likely interacts with the maximal areal mat extent (i.e. by driving resource availability through shading and hydrodynamic disruption), which may be further interrelated with experienced predation pressure. In one instance during observations of fish foraging behavior associated with mats, we observed 494 bites taken on a single relatively unprotected (minimal surrounding physical structure) large-in-area benthic cyanobacterial mat during a time span of 26:05, dominated by bites from *Sc. iseri* individuals (255 and 108 bites from initial phase and terminal phase individuals respectively). This ‘foraging frenzy’ significantly disrupted a majority of the visible mat structural matrix. Interestingly, structural context has previously been hypothesized to mediate herbivorous pressure from reef fishes on basal trophic resources, though with increased herbivorous pressure with greater physical structure (Vergés et al. 2011). This implies that the spatial scale at which these relationships are measured (i.e. seascape vs. local) matter when relating physical context to predation risk. We suggest future research efforts focus on understanding how seascape-scale physical context contributes to heterogeneity in predation pressure experienced by cyanobacterial mat communities and overall temporal asynchrony in community dynamics.

**SUPPORTING FIGURES:**

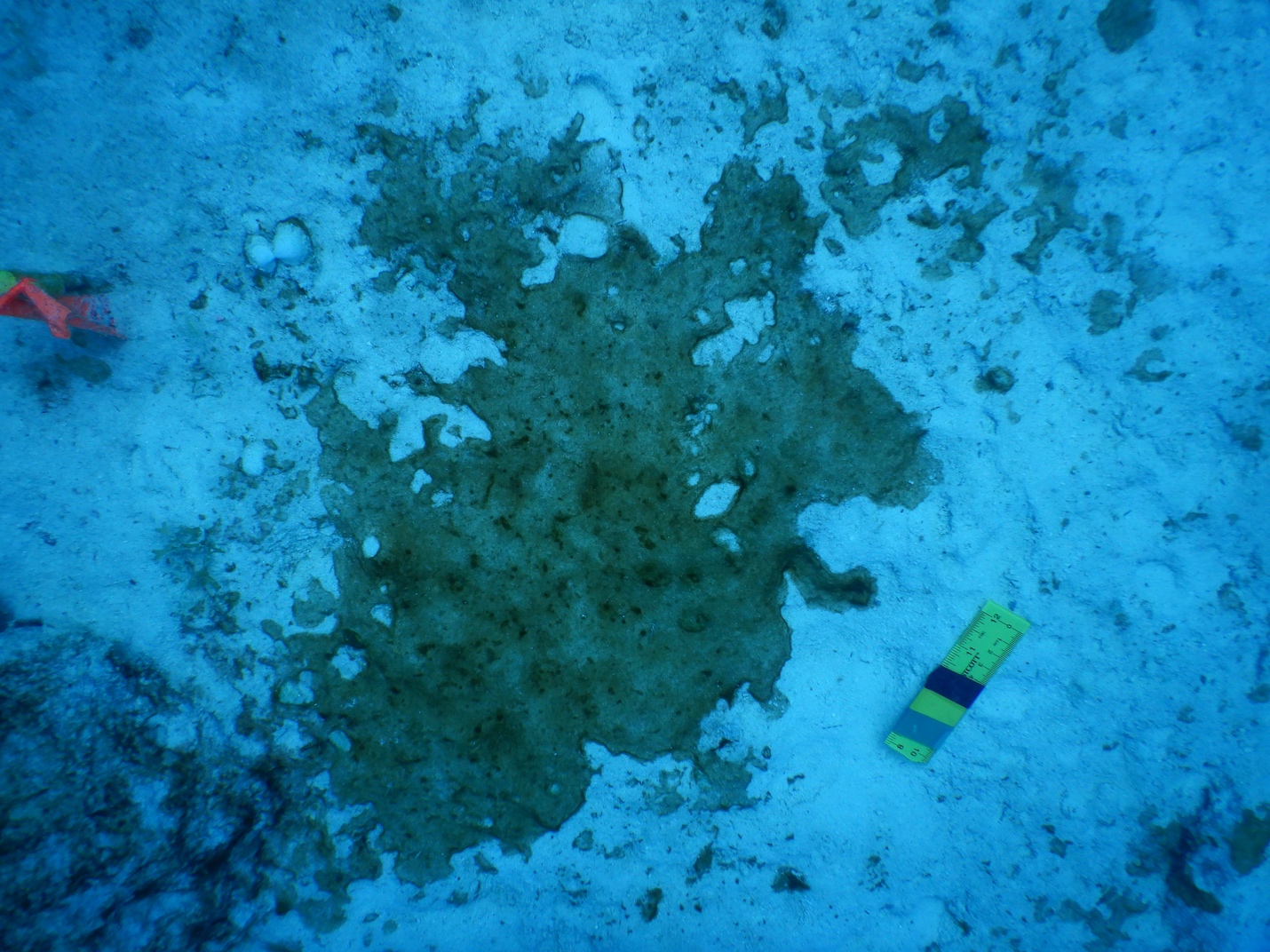

**Figure S1 |** Representative photograph of dominant orange-red cyanobacterial mat morphotype targeted during this study. Image was taken on July 10, 2019 at 09:51AM AST at a depth of 16.3m.

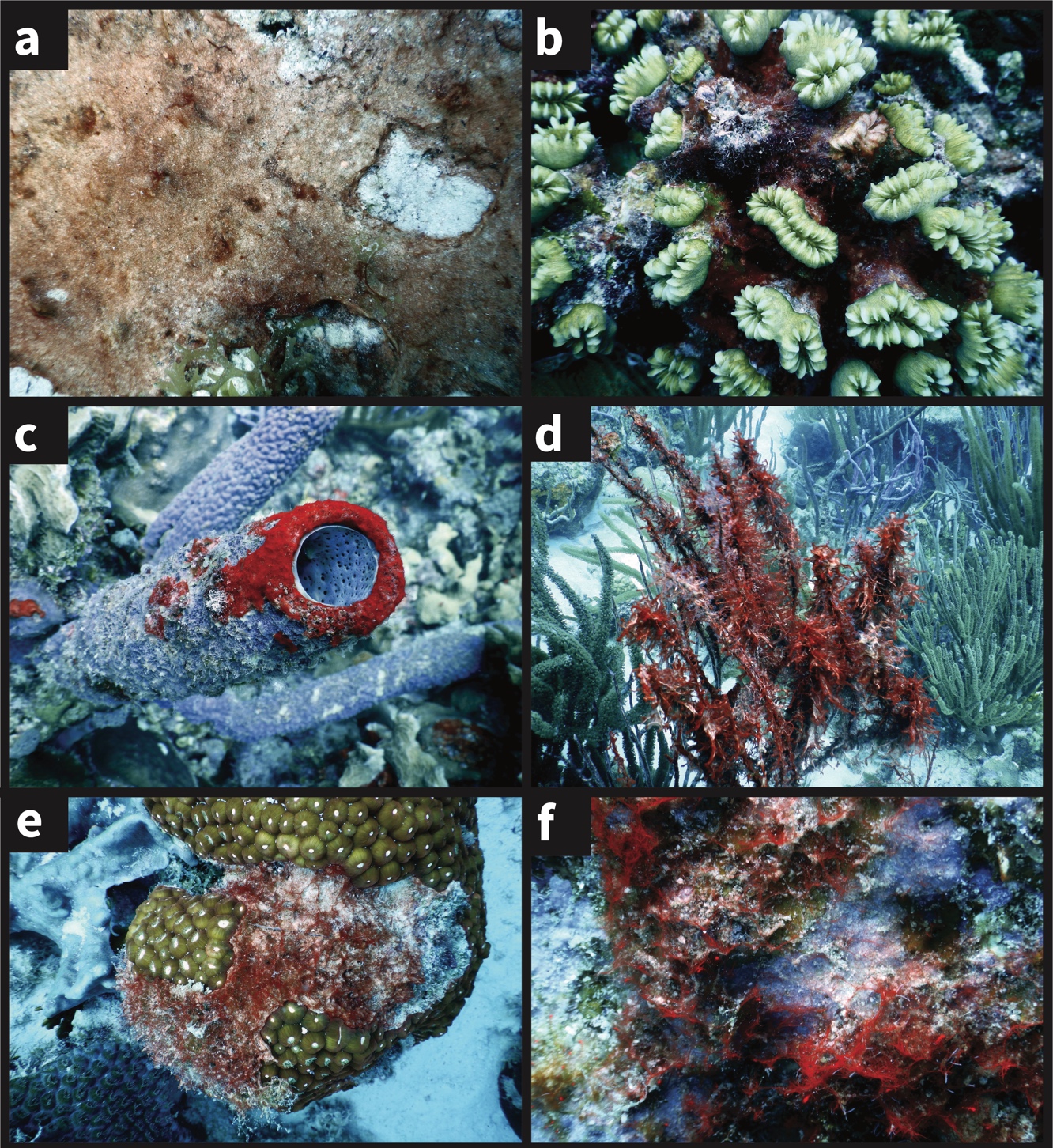

**Figure S2 |** Cyanobacterial mats can be found overgrowing many distinct benthic substrata, including a, Sediment; b, Stony coral; c, Sponge; d, Octocoral; e, hard substrate/EAM; f, Crustose Coralline Algae. All photos by E.C. Cissell in Bonaire, Caribbean Netherlands.

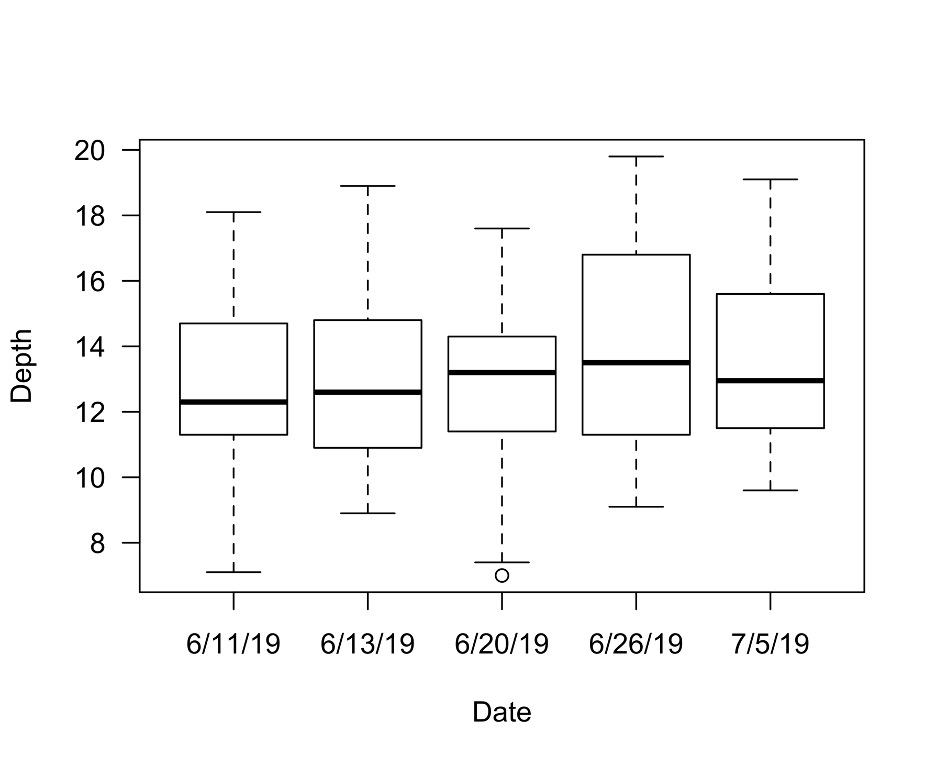

**Figure S3 |** Boxplots of distribution of depth of photoquadrats across sampling dates for metacommunity abundance tracking.

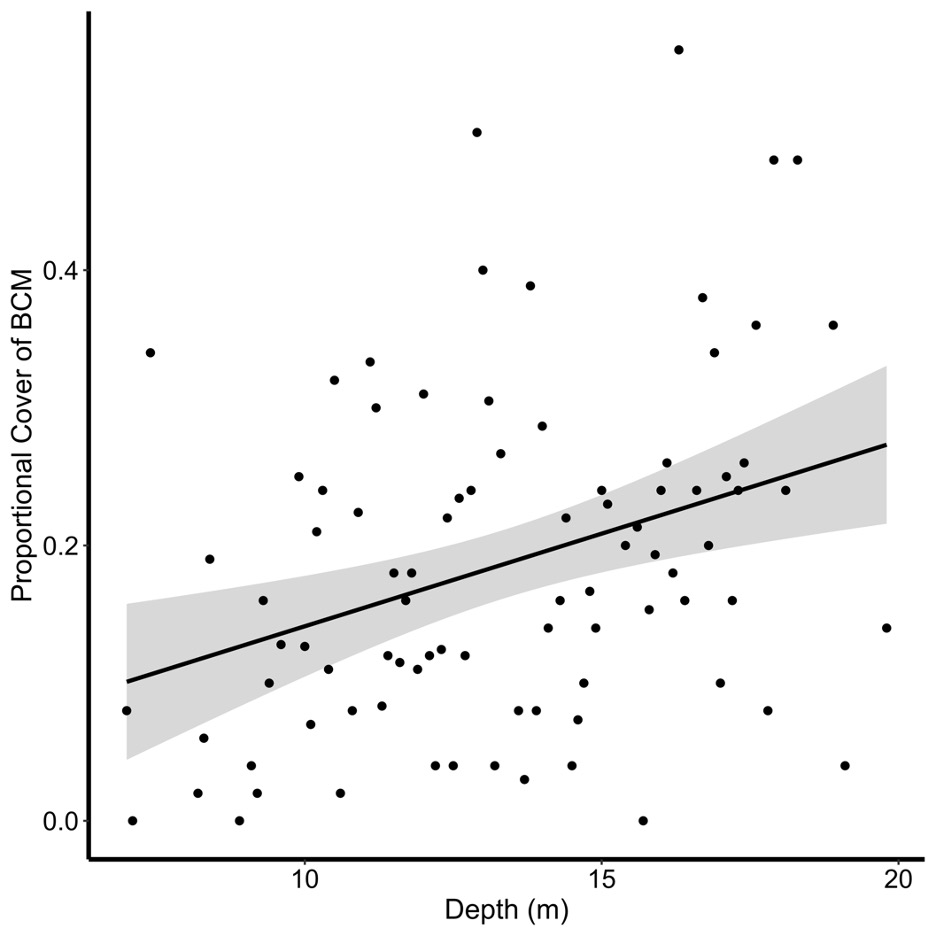

**Figure S4 |** Relationship between site scale proportional cover of cyanobacterial mat and depth of photoquadrat. Positive relationship was significant (LM; p=0.002).

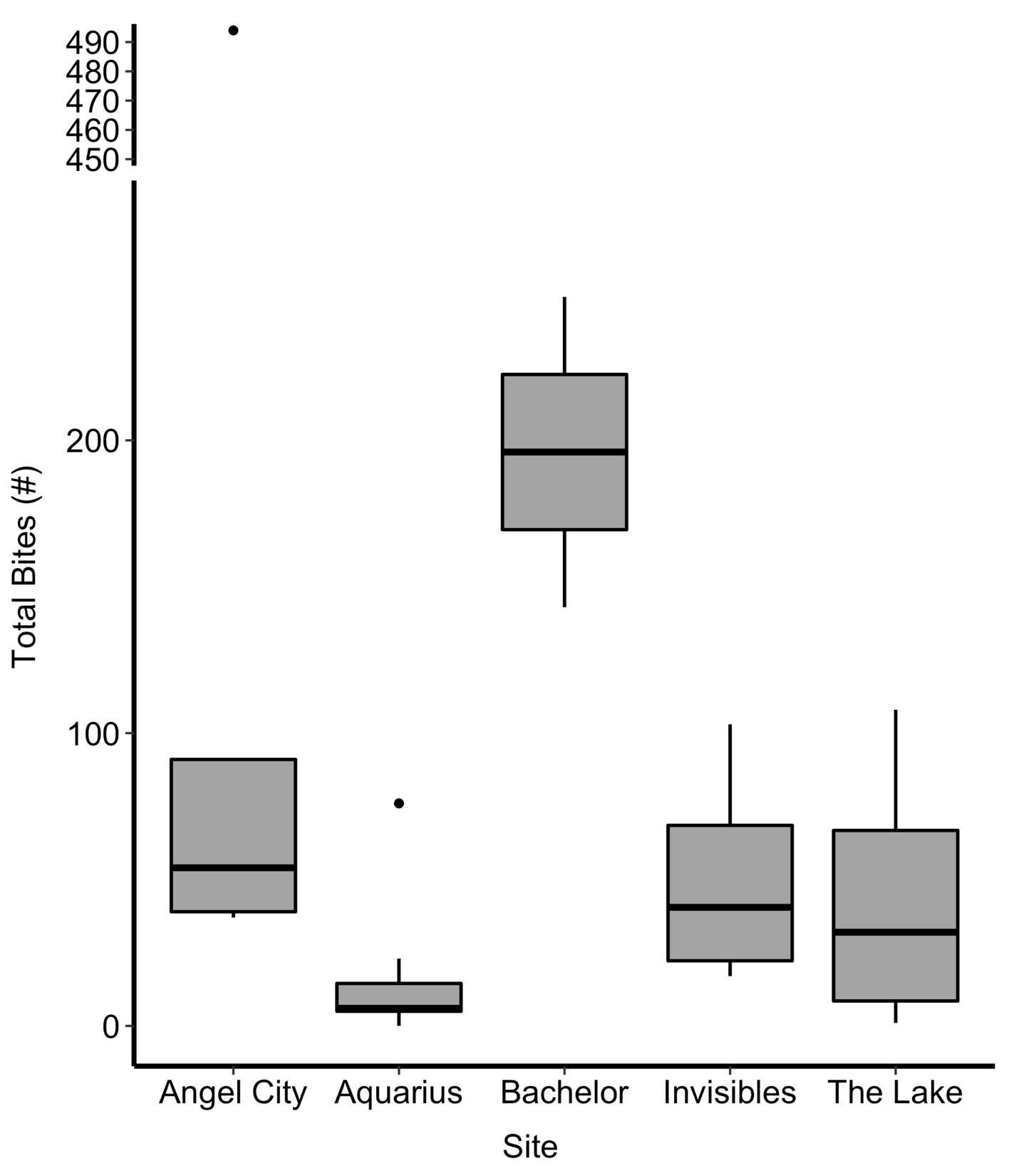

**Figure S5 |** Boxplots of bite count summed across all fish species per sampled site. Note y-axis break during interpretation.

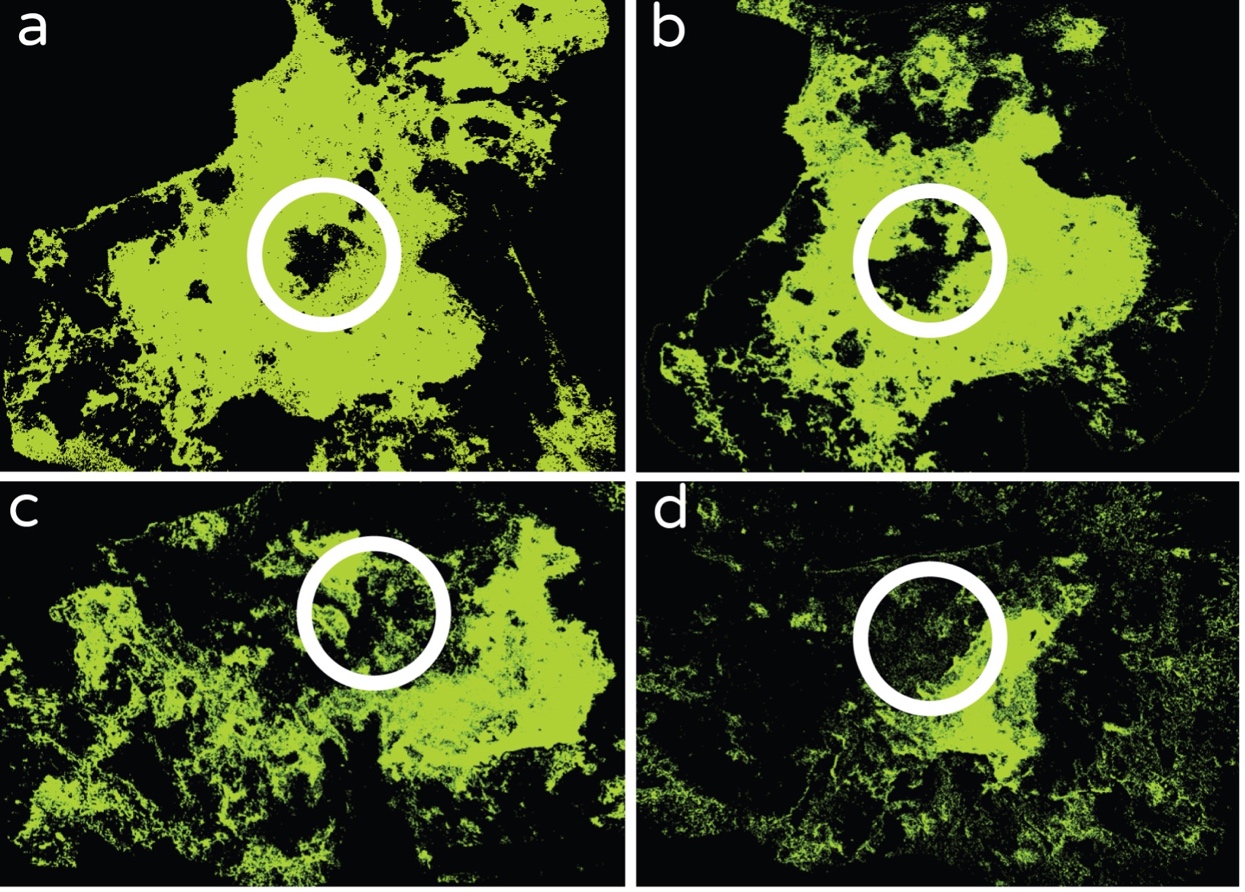

**Figure S6 |** Rasterized cyanobacterial mat showing change in area of a focal cyanobacterial mat (green) across 8 days (a, Day1; b, Day 3; c, Day 6; d, Day 8) with putative grazing scar location circled in white.

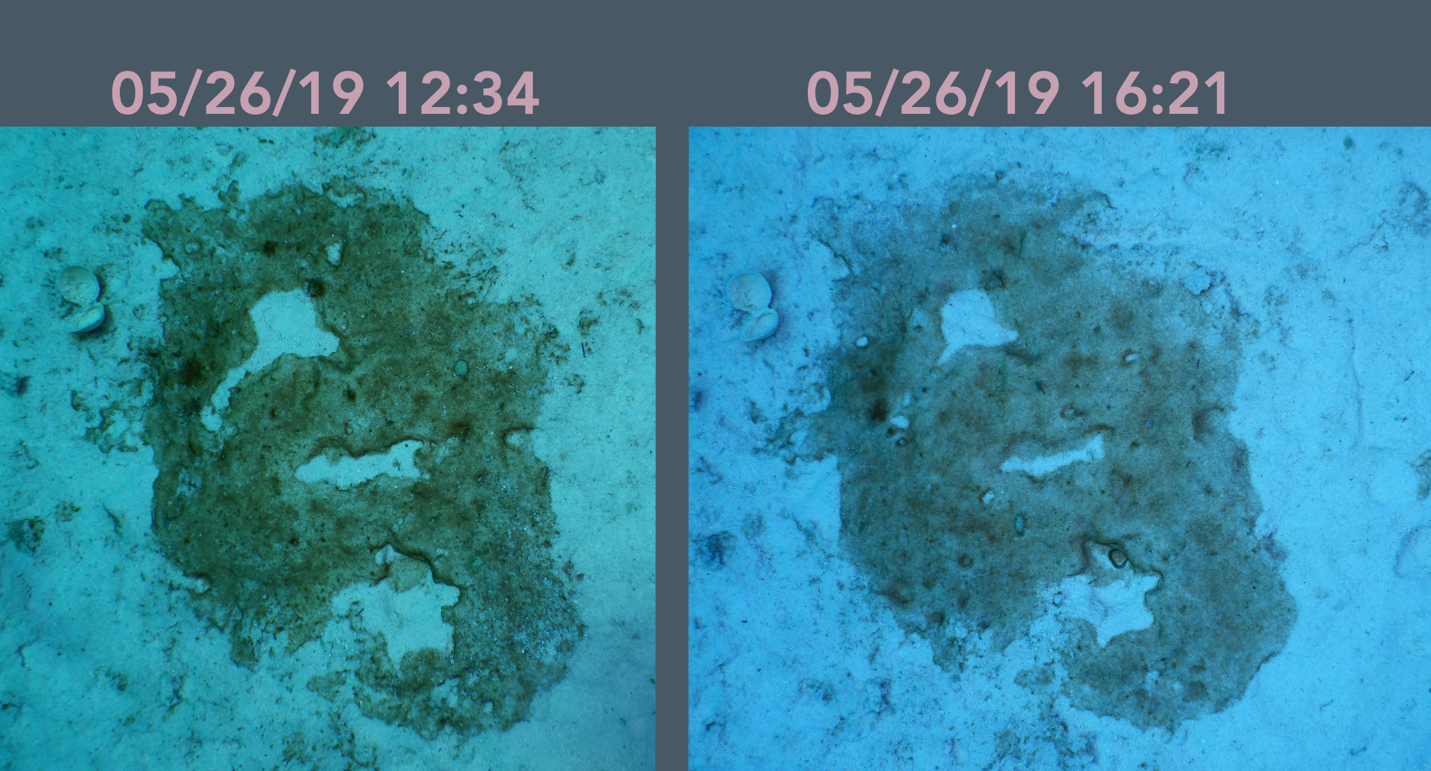

**Figure S7 |** Longitudinal photographs of an individual cyanobacterial mat community collected ~4 hours apart showing rapid colonization of bare space created within mats (circles), which may create mosaicism in the intra-mat landscape of taxonomic and functional richness.

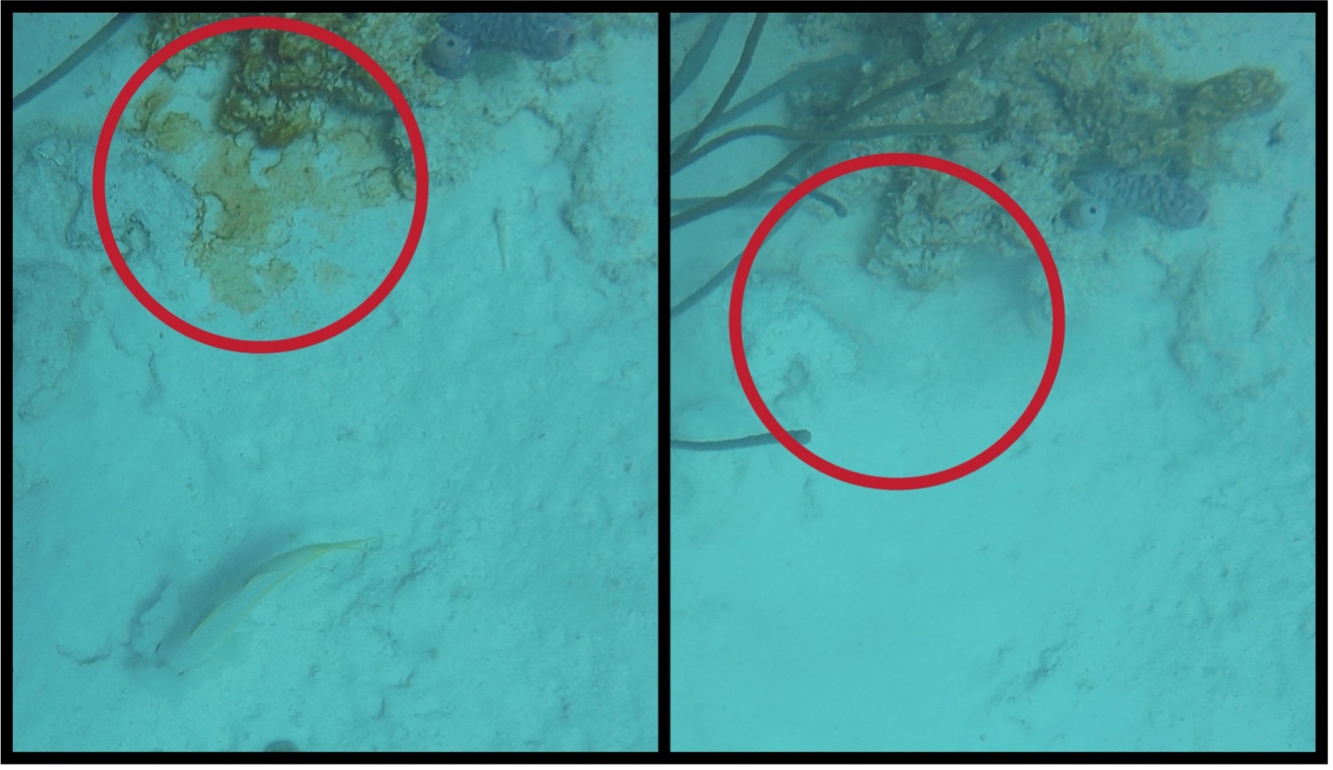

**Figure S8 |** Foraging behavior of the yellow goatfish can indirectly influence cyanobacterial mats, even when foraging is not targeted directly under mat biomass. Shown is a cyanobacterial mat (circled) before (left) and after (right) being completely sedimented following nearby foraging behavior of a yellow goatfish individual (seen bottom of left image).

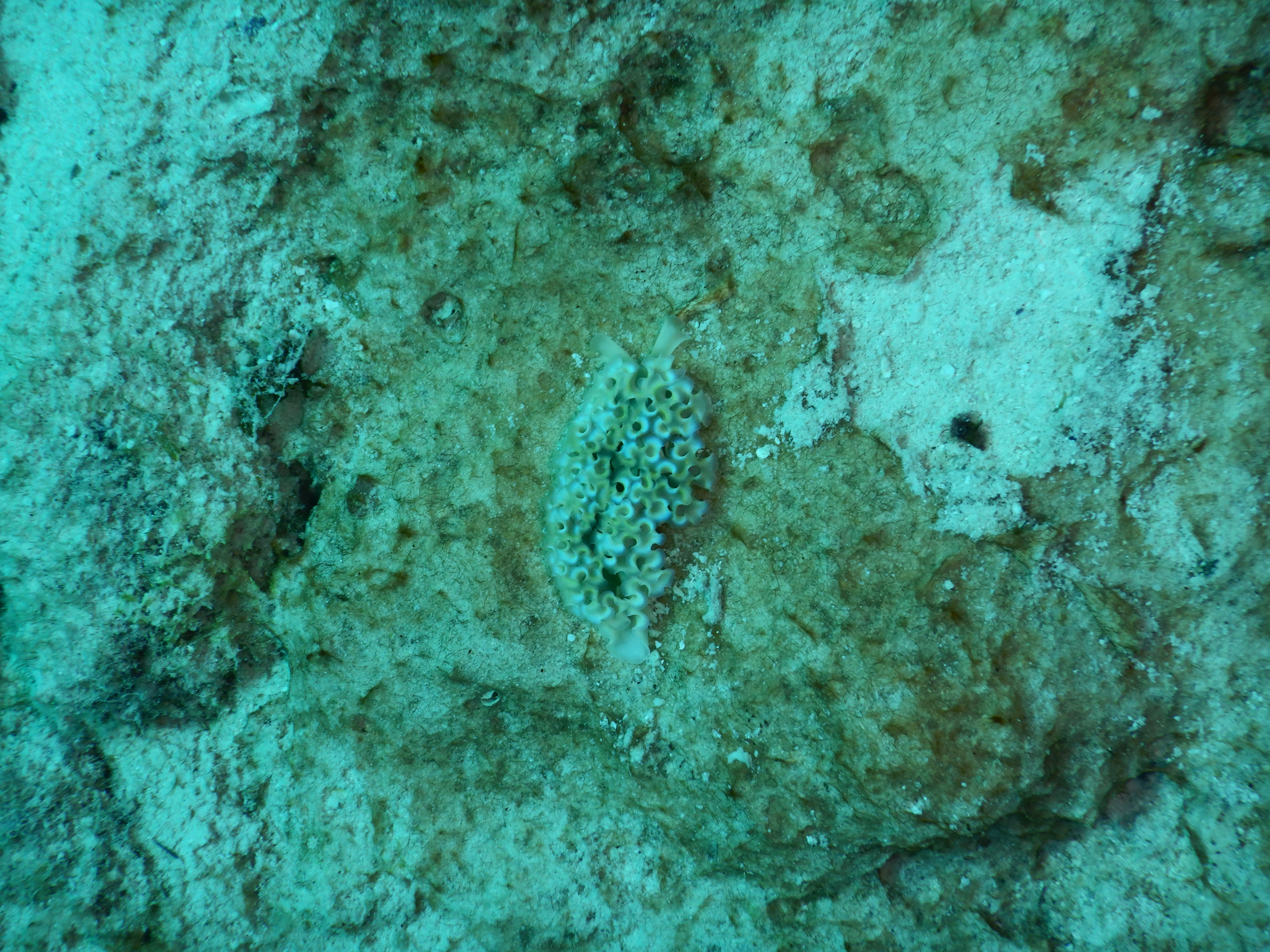

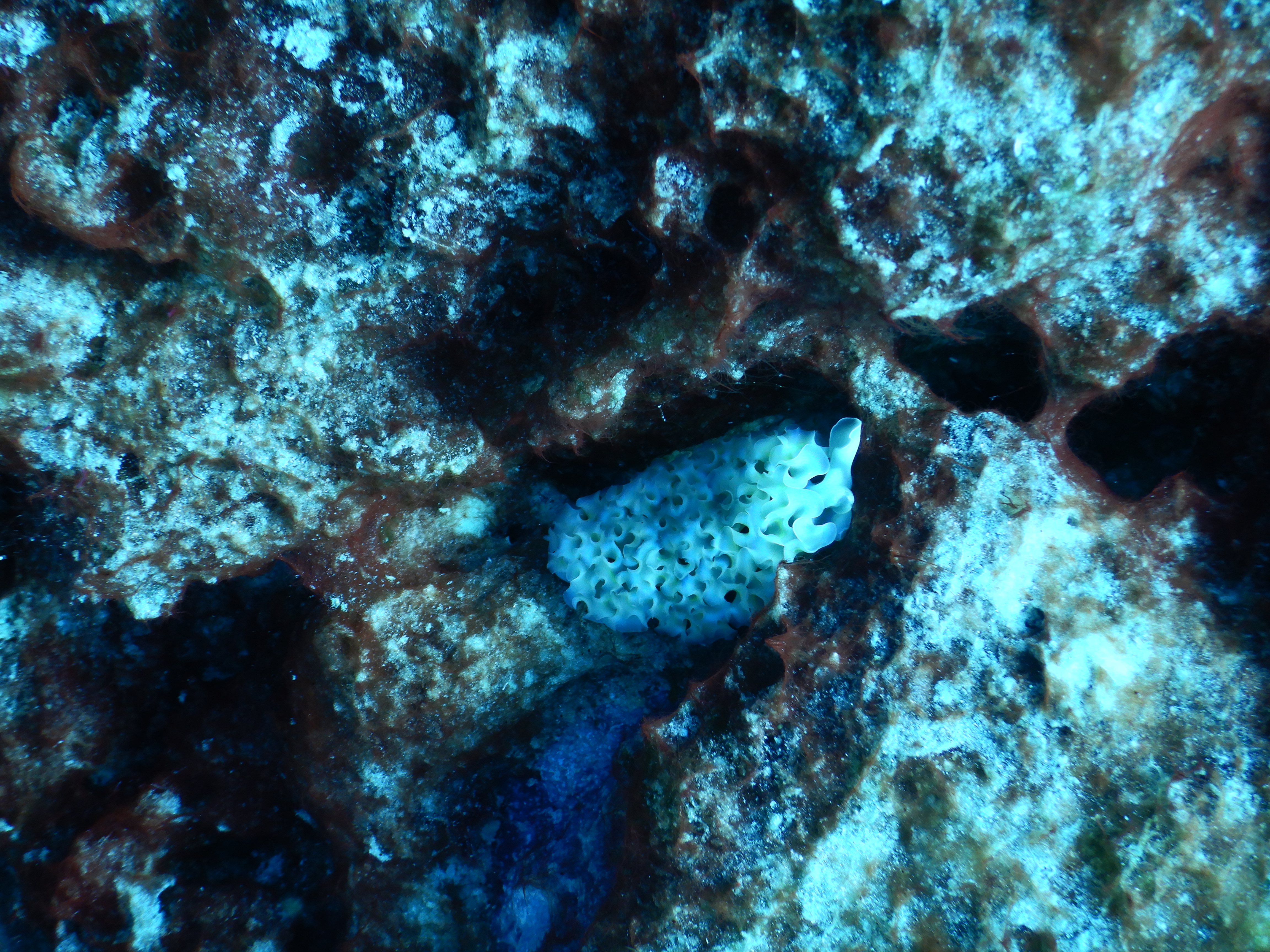

**Figure S9 |** Example photographs of Lettuce Sea Slug individuals observed on sediment- (top) and hard-substrate- (bottom) bound cyanobacterial mats.

**Table S1 |** Numerical probabilities assigned to each varied model parameter for each respective broad grouping (Level).

| **Level** | **DISTURB**  **Goatfish** | **DISTURB**  **Fishes** | **DISTURB**  **Viruses** | **GROWTH RATE** | **DISPERSAL** |
| --- | --- | --- | --- | --- | --- |
| Zero | 0 | 0 | 0 | 0 | 0 |
| Low  Medium | 0.01  0.21 | 0.08  0.28 | 0.01  0.21 | 0.2  0.5 | 0.0001  0.001 |
| High | 0.41 | 0.48 | 0.41 | 0.8 | 0.01 |
